# Ecological stoichiometry and life history theory, not the identity of genomic variants, predict rapid adaptation

**DOI:** 10.1101/2025.11.11.687836

**Authors:** Seth M. Rudman, Joshua J. Faber-Hammond, Ryan E. Sherman, René S. Shahmohamadloo, Sharon I. Greenblum, Heath A. MacMillan, Punidan D. Jeyasingh, Paul S. Schmidt

**Affiliations:** School of Biological Sciences, Washington State University, Vancouver, WA, USA; Department of Biology, Oklahoma State University, Stillwater, OK, USA; Department of Energy Joint Genome Institute, Lawrence Berkeley National Laboratory, Berkeley, CA, USA; Department of Biology and Institute of Biochemistry, Carleton University, Ottawa, ON, Canada; Department of Biology, University of Pennsylvania, Philadelphia, PA, USA

**Keywords:** Predicting Evolution, Ionomics, Experimental Evolution, Evolutionary Ecology, Molecular Evolution, Adaptive Tracking, Population Genomics, Elemental Biology

## Abstract

**Significance:** Forecasting how populations adapt to changing environments is a central challenge in evolutionary biology. We tested the predictability of rapid adaptation by tracking both phenotypic and genomic trajectories of evolution in replicate outdoor populations of *Drosophila melanogaster* and comparing these data to predictions derived from life history theory, ecological stoichiometry, and genomic data. We found that life history theory and ecological stoichiometry forecasted the direction of phenotypic evolution, even with temporal changes in the direction of evolution. In contrast, genomic predictors were largely unable to forecast the SNPs or genes underlying adaptation. These results demonstrate the utility of ecological frameworks to provide general, but robust, forecasts of adaptation in complex environments.

Determining the scale and scope of the predictability of evolution is fundamental to understanding biological processes and to managing biodiversity. Ecological frameworks, including life history theory and ecological stoichiometry, offer testable predictions about the direction of adaptation and trade-offs between traits in response to environmental change. Similarly, well-characterized molecular pathways, genotype–phenotype linkages, and prior evolutionary genomic studies could be predictive of the genomic architecture underlying adaptation. We tested whether ecological frameworks and evolutionary genomic data can be used to forecast rapid adaptation in replicated outdoor populations of *Drosophila melanogaster* evolving in response to natural seasonal fluctuations from summer to late fall. Life history theory predicted the observed pattern of adaptive tracking: in summer, reproductive output increased and stress tolerance decreased, while in fall, this direction reversed, with evolution of increased stress tolerance at the expense of reproduction. Stoichiometric phenotypic evolution was also predictable, with phosphorus and magnesium content, both linked to growth rate, and alkali metal, associated with maintaining homeostasis in response to thermal stress, showing rapid and parallel evolution indicative of adaptation. Temporal genomic data revealed a complex genomic architecture of temporal adaptation and the SNPs and genes involved in adaptation were largely unpredictable. These results demonstrate that ecological frameworks, more than genomic data, have utility in forecasting adaptation in complex and variable environments.

## Introduction

Understanding how adaptation proceeds via natural selection is a fundamental goal of evolutionary biology (1). Whether it is possible to translate understanding of adaptation to prediction—defined as the ability to forecast changes in trait values or allele frequencies from existing information (2)—has long been debated (3, 4). Predicting evolution is critical for managing biodiversity (5–7) and for many aspects of human health and well-being (8–10). Accelerating anthropogenic change, including faster cycling of biogeochemicals (11), threatens to disrupt historical patterns of adaptation and exacerbate phenotype–environment mismatches (12). This heightens the need to translate knowledge of evolution to predicting the direction and pace of adaptive evolution in natural populations (2, 13, 14).

Underlying complexity or contingency could render accurate evolutionary predictions either intractable or impossible (15–17). Genetic drift is well recognized as a process that produces unpredictable evolutionary outcomes (Lande 1975), particularly at modest effective population sizes. Beyond drift, genetic factors including possible constraints (18, 19) or a potential lack of additive genetic variation on which selection could act (20) could likewise reduce predictability. Uncertainty and variability in the ecological factors that drive natural selection may be an even greater source of contingency (21). Despite the many reasons evolution may be unpredictable, there are notable cases of rapid and repeated adaptation, many of which occurred in response to experimentally manipulated agents that are chosen to drive strong directional selection (22–25). Tests of the predictability of rapid evolution in response to natural environmental variation are still comparatively rare and few studies have tested the ability to forecast evolution with empirical data (2). Determining whether existing information, including theory, prior experiments, and environmental data, allows for the identification of both the characters that will respond to selection and the direction of evolutionary trajectories is critical to resolving the extent to which rapid adaptation may be predictable and whether any predictions can be used for applications (4, 26, 27).

Ecological theories could be extended to generate predictions regarding the direction of rapid phenotypic evolution in response to environmental change (28). Life history theory, which focuses on growth, reproduction, and somatic maintenance, has been successful in describing patterns of evolution across space in response to directional selection, including relative investment in reproductive output and stress tolerance (29, 30). Ecological stoichiometry, which considers the balance of energy and materials in ecological interactions and processes (31), has proved tracking flows of elements between organisms and environments can be predictive of ecological outcomes.

Specifically, the growth rate hypothesis posits C-to-P and N-to-P ratios are central limitations to organismal growth rates due, in part, to the high content of P in RNA required for rapid growth (32, 33). Recent work has moved beyond C, N, and P to include the importance of other elements in growth (e.g., Mg (34, 35)) and identifying elemental limitations for specific taxa that can be predictive of their physiology and demography (36–38). This includes elemental contents that are critical to core aspects of life history, including ectotherm responses to thermal stress, where metals are central to maintaining ionic homeostasis (39). Whether elemental contents or the biogeochemical niche more broadly (40) can evolve rapidly in response to natural selection, outside of scenarios of elemental limitation, is largely unknown (41–43) and predictions from ecological stoichiometry have not been used to forecast evolution (38). Together, life history theory and ecological stoichiometry provide testable predictions for the phenotypes, and their directions of change, involved in rapid adaptation.

Predicting evolution at the molecular level is also a major goal in biology (15, 23) and patterns of repeated selection on the same genetic variants illustrate that the genomic architecture of adaptation could be predictable, likely due to underlying constraint or canalization (44–46). Repeated patterns of genetic change have given rise to the hypothesis that the presence of large-effect variants in specific genes (46, 47) or particular SNPs are important for the pace of rapid adaptation and, ultimately, the persistence of populations (13). An improved understanding of whether and how the genetic basis of rapid adaptation is predictable is critical to a range of applications that seek to manage rapid adaptation, from conservation biology to resistance management (48). The polygenicity of many complex traits (49), genetic linkage (23, 50), genotype-by-environment interactions (51), epistatic interactions (52), pleiotropy (53), and genetic redundancy (54, 55) may ultimately limit the ability to forecast the genomic variants that underlie adaptation. Determining overall predictability is critical to understanding the applications of evolutionary genomic data for populations, but few such tests have been done and existing work largely has focused on the evolution of Mendelian traits (22, 45, 56, 57).

Predictions about the genomic architecture of adaptation at the locus or gene level could be made using prior studies or field patterns identifying the genetic architecture of adaptation to similar stressors, genotype–phenotype relationships ascertained through genetic mapping of traits demonstrated to be under selection, or from knowledge of the molecular pathways that shape phenotypes shown to be under selection. All of these sources of genomic information are available to various degrees in model taxa and, when paired with an ‘evolve-and-resequence’ experiment tracking genomic evolution, could be used to test the predictability of the genomic architecture of adaptation (58, 59).

To test the predictability of phenotypic and genomic adaptation, we conducted a replicated outdoor experiment that tracked evolution of eight *Drosophila melanogaster* populations from summer to late fall. We used life history theory and ecological stoichiometry to predict the directions and trade-offs of phenotypic evolution across temporally changing environmental conditions. Specifically, we tested for rapid evolution and temporal trade-offs in life history traits including reproductive output, which we predicted would show evolved increases in summer and subsequent evolved declines in fall. Conversely, we predicted that stress tolerance traits would exhibit tradeoffs with reproductive output and would be most critical in response to seasonally cooling temperatures (60–62), and hence would exhibit declines in summer and evolved increases in fall. We used the framework of ecological stoichiometry to make and test predictions that C:P and N:P ratios would initially decrease, with selection for faster growth and reproduction, than subsequently increase during fall as selection would act to increase somatic maintenance at a cost of reducing growth rates (43). We predicted Alkali metal content (K and Na), which is central to responses to prolonged cold exposure (63, 64), would exhibit increased rates of temporal evolution particularly during periods of seasonal cooling to prevent loss of ionic homeostasis (65). Finally, we used knowledge of molecular pathways controlling ion transport in insects, genotype–phenotype linkages from association studies, and data from prior field evolution experiments to determine whether the genomic architecture of adaptation is predictable. We tested these predictions by assessing parallel evolutionary change indicative of adaptation (66) across eight replicate populations in key life history traits focused on: reproductive output and stress tolerance, stoichiometric basis of growth and thermal tolerance, and temporal whole genome sequencing of pools of individuals from each population (Figure 1).

**Figure 1.**
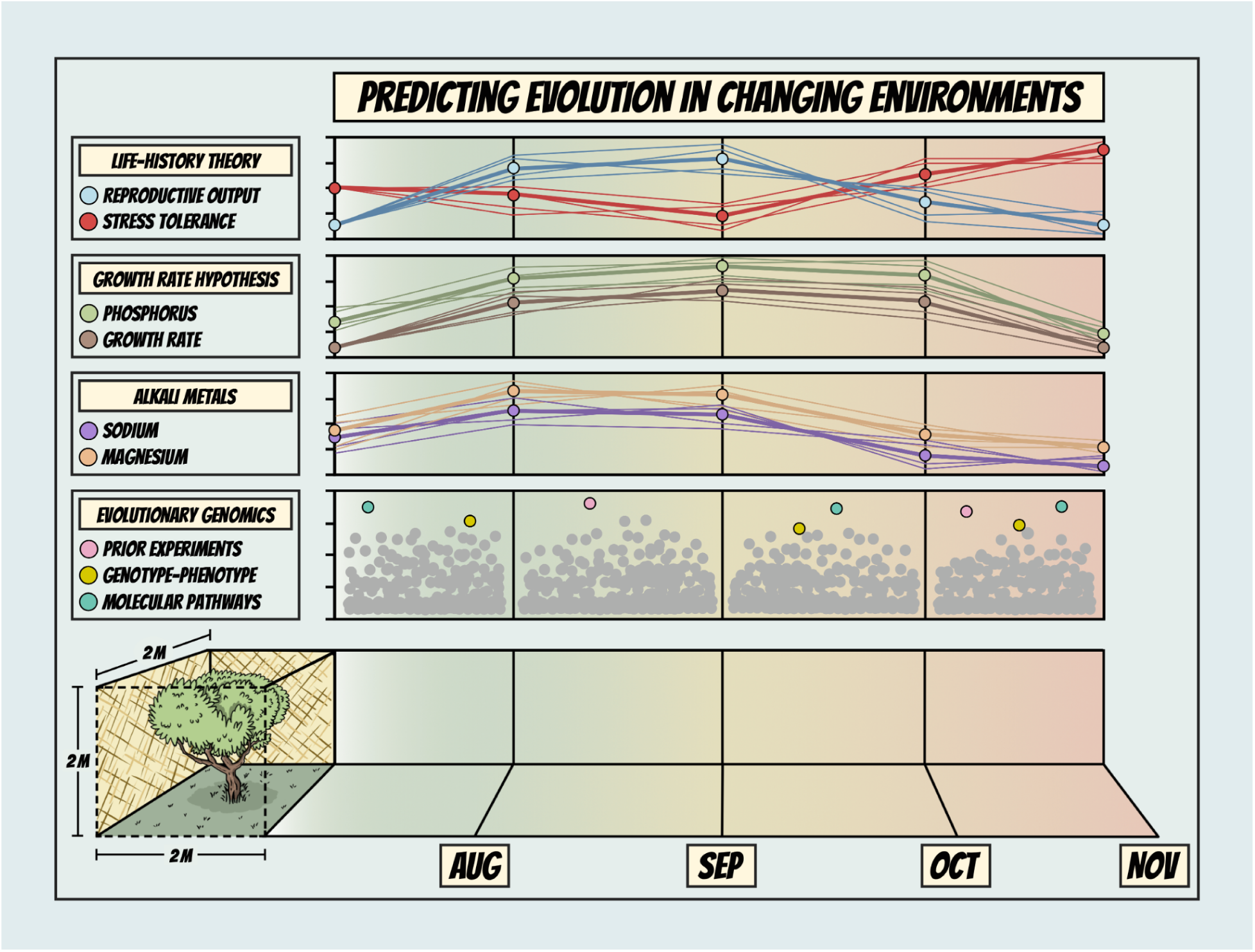
A schematic showing the design and predictions. Tracking temporal evolution of replicate experimental field populations of *Drosophila melanogaster* to test whether adaptation was predictable using: (i) life-history theory (e.g., traits related to reproductive output and stress tolerance), (ii) ecological stoichiometry (i.e., growth rate hypothesis), (iii) major electrolytes (e.g., Na, Mg), and (iv) the evolutionary genomic predictors (i.e., genomic data demonstrating allele frequencies changes in prior field evolution experiments, genotype–phenotype linkages from association studies, and molecular pathways known to influence ion transport). Lines and points shown were created based on predictions derived from theory and projected abiotic change across seasons.

## Results

### Evidence that life history theory predicts rapid adaptation

Temporal common garden rearing revealed parallel phenotypic evolution across independent populations over time, with changes in the direction of evolution indicative of adaptive tracking (Figure 2). Moreover, life history theory, combined with seasonality (Figure S1), largely predicted the direction of evolution in traits related to reproductive output and stress tolerance: selection toward higher output and reduced stress tolerance in summer and lower output and increased stress tolerance in fall. Summer selection between July and August reduced starvation tolerance by >40%, but this was followed by a near doubling of starvation tolerance from September to November (Figure 2A; Chisq = 252.29, *p* < 0.0001). Heat tolerance showed a similar overall evolutionary trajectory, including declines during the hottest months, suggesting the assay may more strongly reflect overall stress tolerance than high heat performance (50% decrease from July to September; subsequent 37% increase from September to November (Figure 2B; Chisq = 83.98, *p* < 0.0001)). All reproductive output traits exhibited patterns opposite to those of stress tolerance traits, with evolved increases in summer and declines in fall. For example, early life fecundity showed an evolved 63% increase from July to October and a 77% decrease from October to November (Figure 2C; Chisq = 92.50, *p* < 0.0001). Similarly, larval development rate evolution showed a 21% increase in rate from July to September and a subsequent 66% slowing from October to November (Figure 2D; Chisq = 1323.4, *p* < 0.0001). Body mass likewise showed evidence of adaptive tracking with a 27% increase in average mass from July to September and a 99% decrease from October to November (Figure 2E; Chisq = 246.2, *p* < 0.0001), fitting with predictions based on the relationship between temperature and fitness (67).

**Figure 2.**
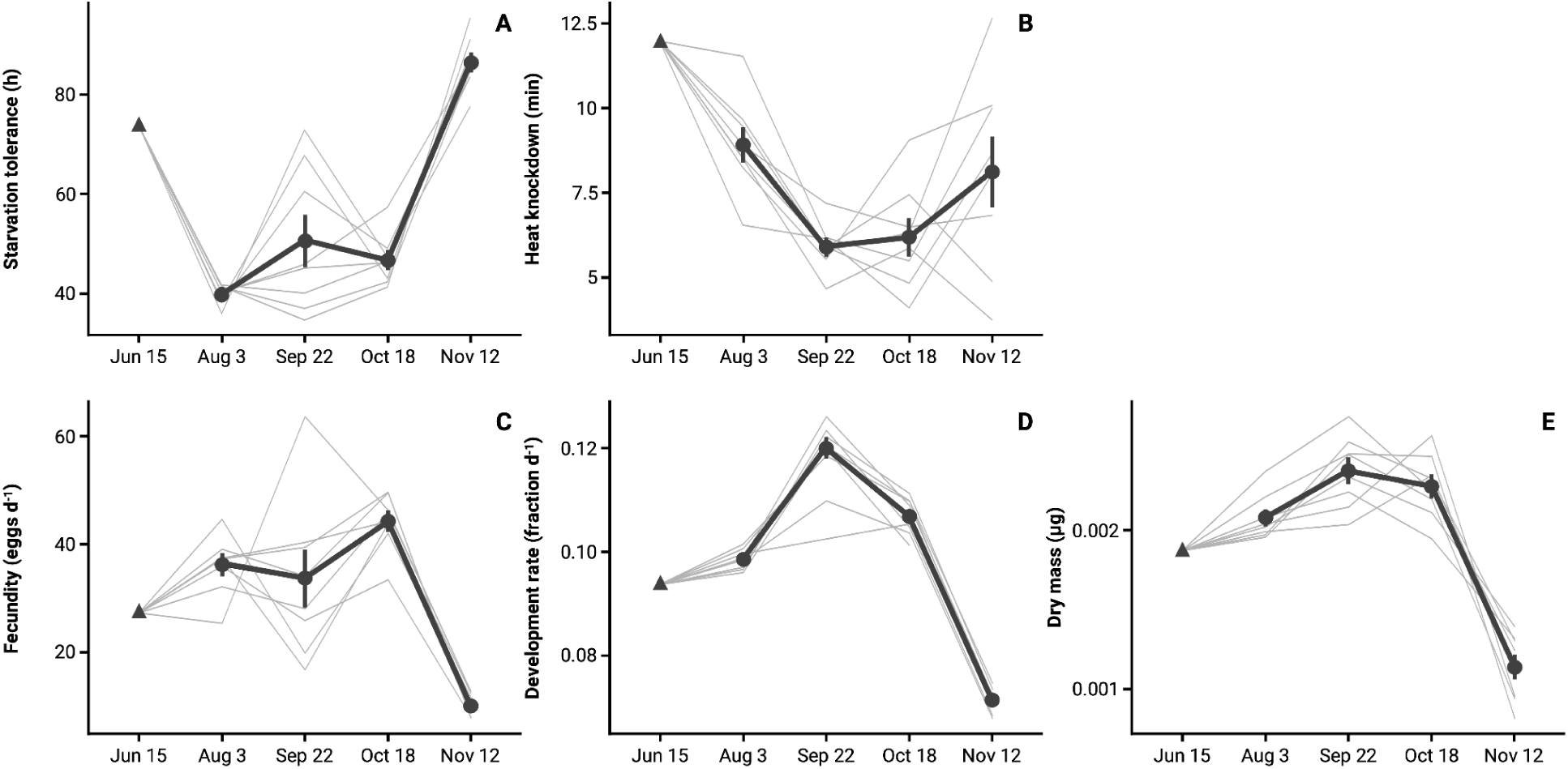
Life history trait evolution over time. All phenotypes were measured after two generations of common garden rearing and hence represent phenotypic evolution. Parallel change over time is evidence of adaptation by natural selection. Panels A (starvation tolerance) and B (heat knockdown) show evolution of a reduction in stress resistance in summer followed by strong evolved increases. Panels C (fecundity), D (development rate), and E (dry mass of pools of five females) show evolution of traits related to reproductive output which show an opposite pattern (evolution of increased output in summer and decline in fall). Gray lines represent individual replicate populations and black points (±SE) and lines represent mean values.

### Evidence that stoichiometry predicts rapid adaptation

The elemental constituents of *Drosophila* (*i.e.,* the ionome (42)) also showed evidence of rapid adaptation with temporal changes in the direction of evolution indicative of adaptive tracking. C:P ratio evolved rapidly (Figure 3A, Chisq = 67.73, *p* < 0.0001) with an initial decrease of 14% from July to August and a subsequent monotonic increase of 99% from August to November, qualitatively fitting with predictions from the growth rate hypothesis. N:P ratio also showed evidence of rapid seasonal adaptation evolution in N:P content across seasons (Figure 3B, Chisq = 53.29, *p* < 0.0001). This included an evolved 54% increase from August to October. Magnesium content, a critical element for stabilizing many ATP dependent processes (35, 68), including growth, also evolved rapidly paralleling patterns observed for P content (Figure 3D, Chisq = 66.84, *p* < 0.0001), with a 24% evolved decrease from October to November. The direction of stoichiometric adaptation, particularly during the fall, fits with predictions for populations with reduced rates of growth and reproduction.

**Figure 3.**
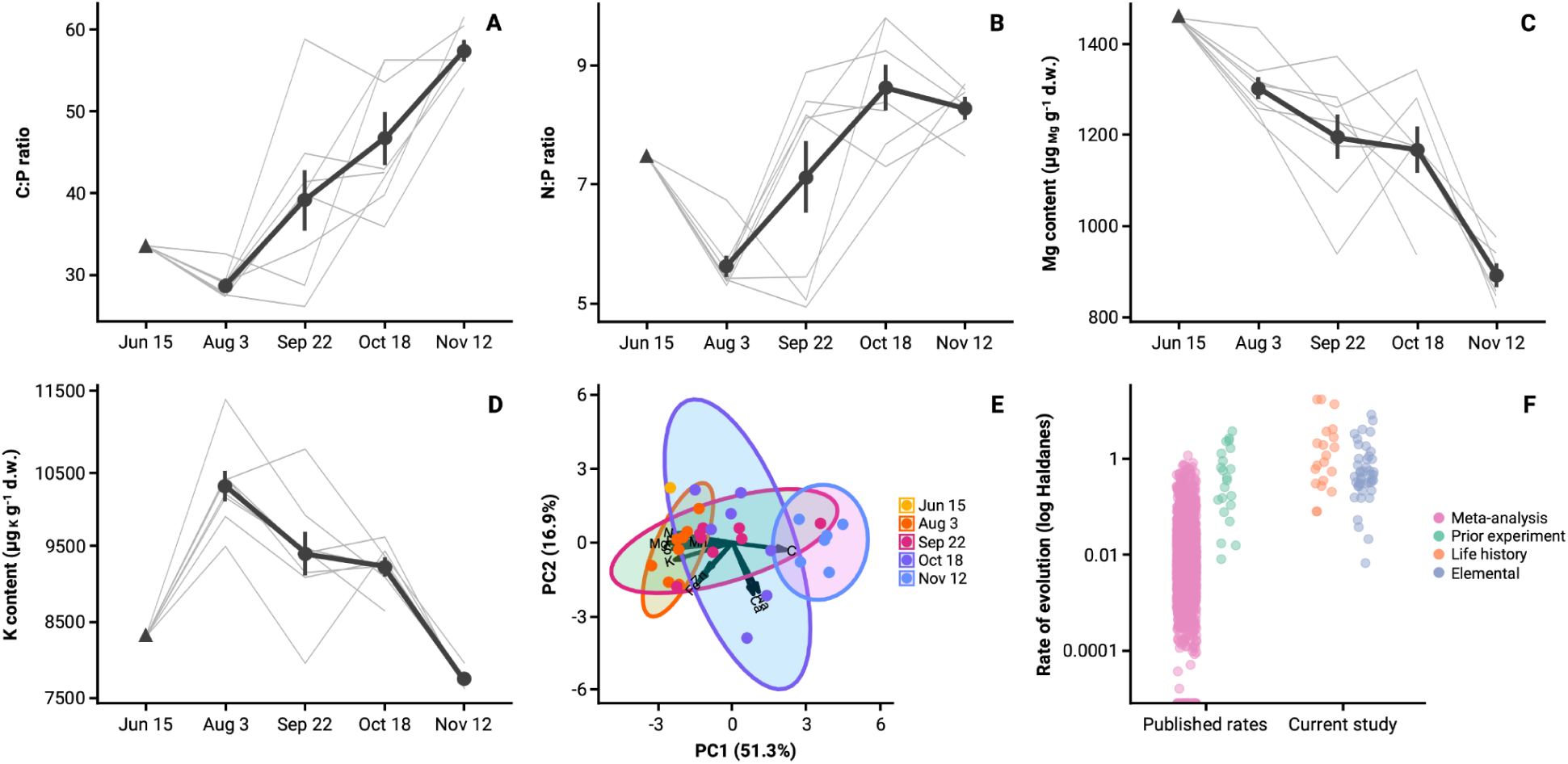
Stoichiometric trait evolution over time. Panels A and B show evolutionary trajectories of elemental ratios key to the growth rate hypothesis: A) The effect of seasonal evolution on C:P ratio; B) The effect of seasonal evolution on N:P ratio. Panel C shows evolution of Mg content, another key element related to growth. Panel D shows patterns of temporal evolution in K, an alkali metal associated with the maintenance of ion homeostasis in cold temperatures in ectotherms. In panels A-D gray lines represent individual replicate populations and black points (±SE) and lines represent mean values. Panel E is a PCA plot showing temporal evolution of replicate populations in 11 biologically active elements (PC1 most strongly influenced by Mg, S, N, K, and P). Panel E shows a comparison of rates of evolution, measured in Haldanes, between prior studies (meta-analysis across tax (71), a prior experiment measuring seasonal evolution of fitness associated phenotypes (57), and the rates of adaptation in life history and stoichiometric traits measured in this study. All phenotypes were measured after two generations of common garden rearing and hence are genetic changes in trait values.

Alkali metal content, crucial for maintaining homeostasis in cold temperatures, exhibited patterns of rapid adaptation. We observed significant and predictable temporal adaptation in K (Figure 3C, Chisq = 96.94, *p* < 0.0001), which included a 24% evolved drop in K content from August to November. Na content largely showed an opposite seasonal pattern with increases during the mid-summer (Supplementary Information, Figure S2, Chisq = 18.48, *p* = 0.001). Overall, we found parallel and rapid evolution of the biogeochemical niche, as principal components based on all elemental data separated populations by time. PC1, which captured 51.3% of the total variation in elemental content, illustrates a clear pattern of temporal ionome adaptation across seasonal time (Figure 3E, Chisq = 21.35, *p* = 0.0003). This parallelism was not observed in the founder populations held indoors during the experiment (Figure S3), reinforcing adaptive tracking as the mechanism driving stoichiometric evolution. Beyond patterns already noted, elements including Fe, S, and Zn showed pronounced patterns of parallel temporal evolution (Supplementary Information, Figure S2), with seasonal declines that coincide with reduced reproduction and shifts in metabolic allocation, fitting with previously described associations (69, 70). Overall, the pace of stoichiometric evolution, measured in haldanes, was comparable to what we observed in life history traits (Figure 4F) and rates of evolution in both classes of phenotypes far exceeded average rates of trait change measured over contemporary timescales (71).

**Figure 4.**
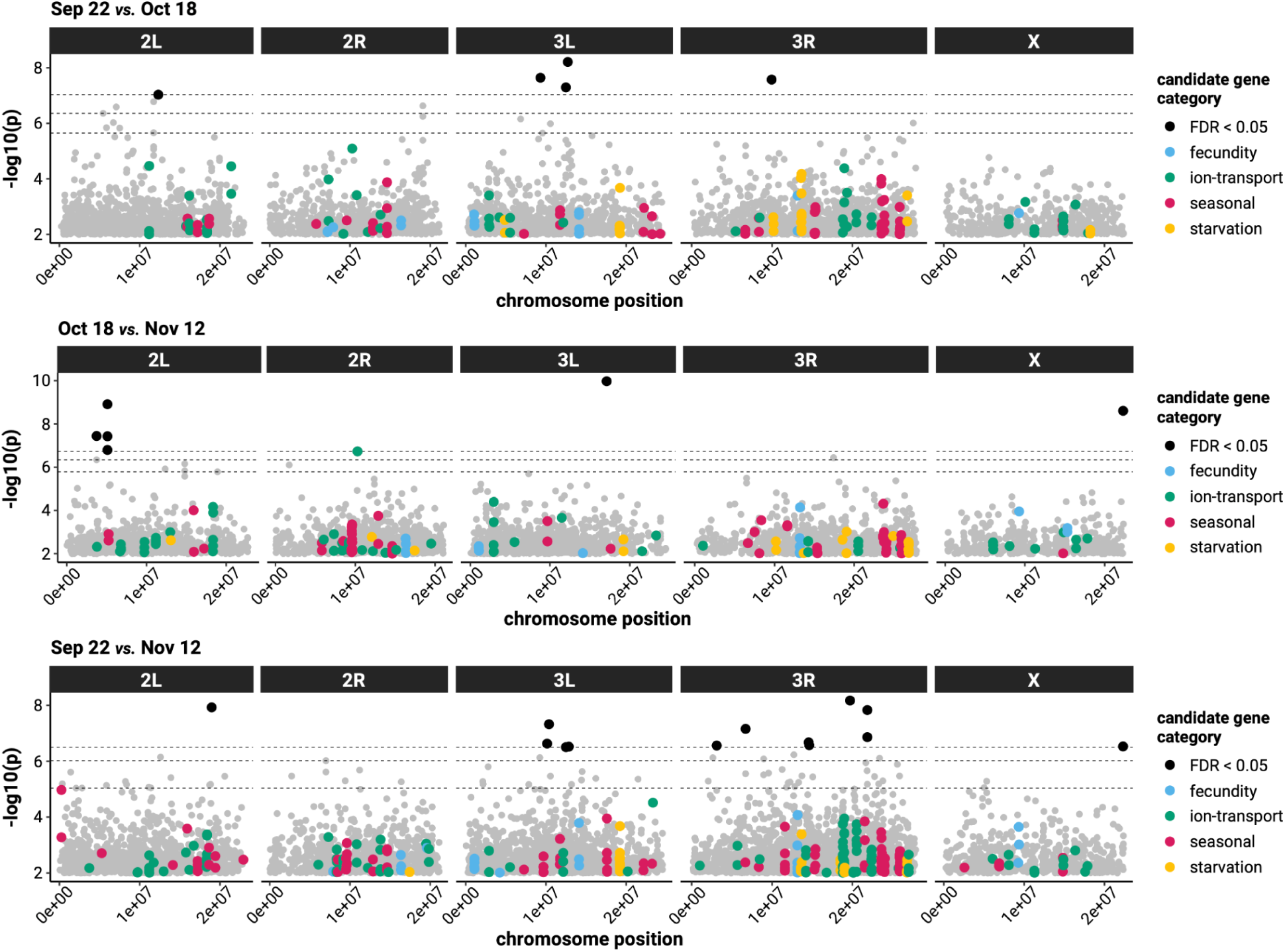
Evidence for adaptation genome wide and at candidate loci. Manhattan plots highlighting SNPs which showed parallel changes in allele frequencies over time with each panel illustrating genomic outliers over a given time interval. Dotted lines from top to bottom correspond to FDR threshold values of 0.05, 0.1, and 0.25, and SNPs with FDR < 0.05 are additionally bolded in black. Sites within candidate genes are colored according to their predictive category (seasonal = prior evolution experiment (57), fecundity and starvation = genotype–phenotypes linkages (73), ion transport = candidate genes based on molecular functions (Table S4)).

### The predictability of the genomic architecture of adaptation

#### Patterns of temporal genomic adaptation, genomic architecture, and gene ontology

In total 13,684 SNPs showed significant parallel allele frequency shifts across replicate populations using an unadjusted-*p* < 0.01 (23 SNPs significant at FDR < 0.05). These putatively adaptive sites were dispersed across chromosomes, with an overabundance on 3R (∼30% of all putatively adaptive sites on 3R which contains ∼23% of total SNPs (Hypergeometric overlap *p* = 1.96E-85)). Known inversions were not enriched for temporal outliers overall, with only In(3R)Mo showing any signature of enrichment (Hypergeometric overlap *p* = 2.90E-3; 5/4173 SNPs on 3R). No significant SNPs at the unadjusted-*p* threshold fell within any other inversions. The potential functions of putatively adaptive sites, as determined through gene ontology enrichment analysis, showed enrichment for largely the same GO terms across all time-point contrasts, suggesting a subset of gene functional categories consistently involved in adaptive tracking (Supplementary Information, Table S1).

#### Tests for predictability of the genomic architecture of adaptation

We used the putatively adaptive loci identified here to test *a priori* hypotheses about the predictability of the genomic architecture of adaptation. Prior experimental work identifying the genomic architecture of seasonal adaptation (57) was not predictive of the adaptive sites we identified here in any of the three experimental time intervals at the SNP or gene level across either significance threshold (Supplementary Information, Tables S2 & S3; Figure 4). Tests for overlap with a larger set of seasonally adaptive SNPs from a multi-year experimental dataset (72) with even greater temporal resolution similarly revealed no hypergeometric enrichment among the top outlier SNP sets significant at the FDR<0.05 level. Genotype–phenotype relationships as identified by GWAS for starvation tolerance and fecundity (73), both of which evolved rapidly in this study (Figure 2), were similarly unable to predict the architecture of adaptation in any contrast at either significance level (Supplementary Information, Tables S2 & S3; Figure 4). Finally, candidate genes known to impact ion-transport (Table S4) did not lend significant predictability for adaptive sites. From a list of 110 candidate gene regions with hypothesized links to ion regulation containing 28,896 SNPs, 62 genes contained 201 SNPs with signatures of selection prior to p-value correction and 1 SNP in the candidate gene *Dh44-R1*, a diuretic hormone receptor with known importance for ion and water balance (74), remained significant after correction (Supplementary Information, Table S2 & S3; Figure 4).

To determine whether overall sets of candidate loci yielded any predictability of the architecture of adaptation we employed Wilcoxon Rank Sum tests to compare overlap between sets of candidate SNPs and both putatively adaptive sites and randomly selected matched sets of loci. When compared to random sets, variants within candidate genes did show some increased overlap with adaptive sites. Most notably, SNPs within ion-transport candidate genes had greater overlap with adaptive sites than random sets for allele frequency change from October to November (*p* = 0.02). Sites from the prior seasonal evolution experiment and loci associated with variation in starvation tolerance showed increased overlap relative to random sets for specific seasonal intervals (Sep-Oct (*p* = 7.1E-04) and Sept-Nov (*p* = 1.0E-02), respectively). In all three cases of enrichment, the average magnitude of allele frequency change in candidate genes was higher than the average allele frequency change observed in the null matched sets. However, the SNPs driving these patterns in seasonal- and starvation-associated genes largely differ from the candidate SNPs themselves and we observed no significant predictability at theSNP level for any set of predictors (Supplementary Information, Tables S5 & S6).

## Discussion

The potential applications for evolutionary forecasts are clear; rapid adaptation has major influences on human health, ecosystem services, and the conservation of biodiversity (9, 13, 75, 76). Patterns of repeated evolution demonstrate that evolutionary outcomes can be shaped by the deterministic force of natural selection (77). This includes work detailing rapid and repeated adaptation of *Drosophila* populations in field studies (62, 78, 79), laboratory experiments (54, 80), and field experiments (57, 72, 81, 82). We likewise found repeatability in evolutionary trajectories across replicate populations, beyond what is often seen when tracking evolution in field eukaryotic populations (77, 83). This included patterns of temporal variation in the direction of adaptation indicative of adaptive tracking, inverse evolutionary trajectories of life history traits related to reproductive output and stress tolerance suggestive of temporal trade-offs, and adaptation in stoichiometric phenotypes not previously known to evolve rapidly in response to natural environmental variation (43, 57, 62). There is little in the biology of *Drosophila* or the specifics of this experiment, in which natural seasonal variation and associated ecological changes are the primary agents of selection, to indicate the evolutionary trajectories we observed should be exceptional amongst the genetically diverse multivoltine taxa that are central to many ecological communities (84, 85).

Here we translate these hallmarks of evolutionary determinism produced by natural selection to test predictions about adaptation derived from ecological frameworks, evolutionary genomics, and genotype–phenotype linkages. The direction of phenotypic evolution largely fits with predictions derived from the ecological frameworks of life history theory and ecological stoichiometry. Both frameworks can be combined with information on environmental change to forecast directions of trait evolution and potential trade-offs (31, 86, 87). Determining the utility of these ecological frameworks for predicting evolution across taxa and timescales is important for ascertaining generality and potential applications of these predictors (88). Ultimately, integrating knowledge of the direction of evolution into ecological niche models and biodiversity forecasting is critical to increasing the accuracy of conservation projections and management (13, 89).

Prior experimental work using *Drosophila* has demonstrated both rapid phenotypic adaptation and that a considerable portion of the genome shows parallel allele frequency changes across replicates due to the combination of genetic linkage and selection (57, 72, 90). While we detect parallel allele frequency change over time, the number of outlier loci and the size of haplotype blocks shaped by selection are considerably smaller than in prior experiments. Selection was strong with rates of phenotypic evolution meeting or exceeding those found in any previous field experiment (Figure 3F), likely due to greater genetic diversity. The most likely causes for the differences in genomic architecture relative to prior experiments are sampling protocol and the underlying genomic composition. We tracked changes in allele frequencies by sequencing a sub-sample of the full population of adults, differing from previous protocols that sequenced pools drawn only from reproducing individuals (57, 72, 90). The experiments differed also in genomic composition, here we used isofemale lines to form a genetically diverse founder population instead of outbreeding previously inbred lines, which have tremendously inflated genetic linkage beyond what is seen in natural *Drosophila* populations (e.g., correlation coefficients between alleles are often <0.1 within <500bp in nature (91)). While selection was strong, it is likely the reduction in genetic linkage, and associated reduction in genetic draft (pseudohitchhiking (92)) (57, 72), led to a far lower proportion of the genome undergoing parallel allele frequency changes (Figure 4). As a consequence, outlier loci identified in this study are much more likely to be the direct targets of selection. Continued investigations into the relationship between genetic linkage, the genomic architecture of adaptation (93, 94), and predictability, including predictions made across genomic backgrounds, will substantially advance our understanding of epistatic constraint (95) and the mechanistic genetic basis of adaptation.

In contrast to the overall predictability derived from the combination of environmental patterns and ecological frameworks the genomic architecture of adaptation was effectively unpredictable. This is a notable departure from prior investigations of the genomic architecture of adaptation in eukaryotes that assessed selection on specific large effect variants, which often do show elevated signatures of selection (22, 45, 47, 56, 96). The genetic architectures of the traits under selection in this experiment are polygenic (73), hence a large number of interactions could increase interference or epistatic effects (50, 97), and selection both acted on multiple traits and varied temporally (Figures 2 and 3). Temporal reversals of dominance could further reduce the strength of genotype–phenotype linkages and limit predictability (98, 99). Moreover, the extent to which predictability is possible across populations or years, where genetic composition varies, is largely unknown (2). Together, these factors may render the genomic architecture of adaptation largely unpredictable in natural populations with polygenic fitness traits (49, 100, 101), even if there are common variants with moderate fitness effects (97). The unpredictability of the genomic architecture of complex trait evolution is emerging as a pattern across studies, including cases of *Drosophila* pigmentation (90) and mouse body size evolution (102). Yet, it is possible that increasing the quality of predictors, including conducting genotype–phenotype mapping in field environments, building extensive temporal datasets (96), or overall improvements in understanding the molecular mechanisms responsible for stress responses, may increase the predictability of the genomic basis of adaptation. Given the effort required to generate these data, the identification of specific genetic variants likely important for future adaptation may only be translatable to evolutionary applications when there are large effect loci that have been identified or verified in a field realistic context. In scenarios where fitness is composed of many traits, each with a polygenic basis, identification of causal loci or genetic variants important for future performance may be impractical (13). The limited, but significant, patterns of predictability we observed at the gene level do reinforce some potential utility of broad gene or pathway level knowledge for predicting adaptation (7). Overall the lack of genomic predictability, and the striking utility of ecological frameworks for evolutionary prediction, reinforces that trait-based, ecological, or quantitative genetic approaches to predicting rapid adaptation are likely more useful than a focus on putatively adaptive alleles. Translating evolutionary predictability into improved forecasting of demography and persistence in response to environmental change is critical to improving the management of biodiversity and ecosystem services.

## Methods

### Seasonal evolution experiment

To begin the temporal evolution experiment we constructed an outbred founding population of *Drosophila melanogaster* by crossing 150 wild-collected isofemale lines that were collected from Pennsylvania in 2016 and 2017. These lines were not inbred and hence each likely retained substantial genetic diversity (103). 10 males and 10 females were taken from each line and combined into a large breeding cage. After three generations of mating and density controlled rearing in lab conditions (25°C; 16L/8D) we introduced 500 females and 500 males into each of eight outdoor experimental cages on June 15th, 2017. Outdoor cages are 2m × 2m × 2m enclosures constructed of fine mesh built around metal frames (see (104) for additional details). Inside of each enclosure we planted one peach tree and vegetative ground cover to provide shading and physically mimic the natural environment.

Peaches were removed before ripening to prevent flies from feeding on them. Each of the eight populations were fed 300 mL of a standard *Drosophila* media three times a week (modified Bloomington recipe) from June 15th to November 12th, 2017.

### Measuring evolution in fitness associated phenotypes

To track phenotypic evolution we collected 1,200 eggs from each experimental replicate at four intervals during the experiment (August 3rd, September 22nd, October 18th, and November 12th). These eggs were reared for two generations in laboratory common garden conditions (25°C; 16:8 light cycle; density controlled) to allow for the measurement of phenotypic evolution by removing any environmental variation. We assessed the extent of evolution in five fitness associated phenotypes related to reproductive output and stress resistance. Stress resistance phenotypes included starvation tolerance (time to starvation for each sex) and heat knockdown (time to knockdown at 55°C). Traits related to reproductive output: fecundity (number of eggs produced per female per day on days 3-6 post hatch), larval development time (1/time of development from egg to adult), and body mass (pools of 5 females). These focal phenotypes are commonly measured, related to organismal and population growth, and associated with major fitness life history trade offs (e.g., stress tolerance and reproductive output) (57, 62).

Following life history theory we predicted that the direction of phenotypic evolution would favor increased rates of development and reproductive output with potential trade-offs of reduced stress tolerance during summer conditions. We predicted the inverse in fall with selection driving increases in stress tolerance with a trade-off towards reduced reproductive output. We tested for adaptation as parallel change across replicate cages over time in each phenotype using linear mixed effects models with time point as a fixed effect and experimental cage as a random effect (‘lmer’ package). To assess significance we used a Wald chisq test (‘car’ package).

### Stoichiometric phenotypic evolution

To test the predictability of rapid adaptation from stoichiometric principles we generated multi-element (‘ionomic’) data from five experimental populations at each of the four sampling periods (August 3rd, September 22nd, October 18th, and November 12th). To do so, flies were sorted into pools of five females and stored dry at −80℃. For each population at each of the four sample periods, we generated elemental composition data from three samples composed of four individuals in each sample. Each sample was dried at 60℃ for ∼72 hrs until they reached a constant dry mass and weighed using a microbalance (Mettler Toledo XP2U). An automated CHN analyzer (varioMicro Cube; Elementar Americas, Mt. Laurel, NJ, USA), was used to determine C and N concentrations for all *Drosophila* samples. To determine the concentrations of the other elements, we used an automated ICP-OES analyzer (iCAP 7400; Thermo Scientific, Waltham, MA, USA). Before analysis, *Drosophila* ICP-OES samples were digested in 15 ml trace metal-free polypropylene conical centrifuge tubes (VWR International, Radnor, PA, USA) using a 2:1 mix of trace metal grade 67-70% HNO_3_ (BDH Aristar ® Plus, VWR International, Radnor, PA, USA) and trace metal grade 30-32% H_2_O_2_ (BDH Aristar ® Ultra, VWR International, Radnor, PA, USA). Each digested sample was diluted with Type 1 ultrapure water to a final concentration of 5% v/v of acid. Validation and calibration of the ICP-OES was achieved by using multi-element external reference standards (CCV Standard 1, CPI International, Santa Rosa, CA, USA). Additionally, an in-line Yttrium internal standard (Peak Performance Inorganic Y Standard, CPI International, Santa Rosa, CA, USA) was used to correct for any instrument drift or matrix effects. Digestion blanks were also run to correct for background concentrations. Concentrations of elements within samples that were within range of the standard deviations for the blank controls were excluded from further analysis, as these values are indicative of concentrations near or below the limit of detection of the ICP-OES. Nine elements (calcium, Ca; iron, Fe; potassium, K; magnesium, Mg; manganese, Mn; sodium, Na; phosphorus, P; sulfur, S; and zinc, Zn) were above detection limits, and the concentrations of these elements along with C and N (in μg/g) were log_10_-transformed before statistical analyses.

Statistical analyses focused on testing predictions derived from the growth rate hypothesis and knowledge of ionoregulatory homeostasis challenges in response to cold temperatures. We tested for temporal evolution in C:P and N:P ratios and Mg content, all central to organismal growth rate, with the hypothesis that these ratios would decrease during periods of selection for increased reproductive output. In addition, we tested for evolution of Alkali metal content, which is critical to the maintenance of organismal homeostasis in ectotherms. A particular focus was testing for decrease in K content in response to seasonal cooling, as reduced content could reduce the onset of hyperkalemia, a major cause of loss of homeostasis and death in cold (39, 64, 65). We used linear mixed effects models with time treated as a fixed effect and population as a random effect test for evolution of each phenotypic and stoichiometric response. To assess significance we used a Wald chisq test (‘car’ package). There is little known about the pace of stoichiometric evolution (Lemmen 2019). To determine whether overall stoichiometric profiles show evidence of adaptation as evidenced by parallel change across replicates we conducted a principal components analysis, extracted values for PC1, and used a linear mixed effects model with time treated as a fixed effect and population as a random effect.

### Genomic sequencing and identification of candidate loci

To assess the genomic basis of rapid evolution in response to seasonality we sequenced pools composed of 120 males and 80 females collected from each outdoor cage on September 22nd, October 18th, and November 12th. In addition, we also sequenced two pools of individuals from the founder population. We extracted DNA and prepared libraries using ∼500bp fragments for whole genome sequencing (KAPA Hyper Prep kit). Libraries were sequenced on Illumina NovaSeq systems with 150bp paired end reads. Reads were checked for quality using FastQC. Adapters were trimmed with Skewer (105) and reads with a quality score <20 were removed, and overlapping read pairs were merged with PEAR (106). We aligned reads to a reference genome composed of the *Drosophila melanogaster* reference sequence (v5 (107)) using BWA (108), then removed duplicate reads with Picardtools and realigned remaining reads around indels with GATK’s IndelRealigner (109). After mapping and QC we retained an average of 83M mapped reads per sample at an average coverage (mosdepth (110)) of 109x of the *Drosophila melanogaster* autosomes (range 92-133×) and average coverage 92× on the X chromosome. We then used PoPoolation2 (111) to obtain allele counts at segregating sites, discarding bases with quality <20. To be included for downstream analysis we required SNPs to: be bi-allelic, not contain an indel in any sample, have an average minor allele frequency >0.05% across all samples, and have at least 50 mapped reads total in every sample. Additionally, we excluded SNPs within repeat regions as defined by UCSC RepeatMasker (112). This yielded a dataset of ∼1.8 million SNPs.

#### Identifying patterns of temporal genomic adaptation

To identify the genomic architecture of adaptation we used generalized linear models (GLM) to test for parallel changes in allele frequency at each SNP over each experimental time interval. GLMs treated time-point as a fixed effect and used a ‘quasipoisson’ error model implemented through the *glm* tool from the stats R package. *P*-values for consecutive time-point and first to last time-point contrasts were extracted via the Emmeans R package. Resulting *p*-values were FDR adjusted with *p.adjust* from the stats r-package using the FDR method (113). To determine whether putative allele frequency differences were associated with structural variation we tested for hypergeometric overlap of significant outlier SNPs with linkage groups and known inversion regions (57). Additionally, we conducted gene ontology enrichment analysis for genes with SNPs showing strong signatures of selection, before and after FDR correction, using the BiNGO v3.0.5, implemented in Cytoscape v3.10.2 (114, 115).

#### Predicting loci under selection from experiments, genotype–phenotype linkages, and molecular pathways

We tested for predictability by assessing the overlap between outlier sites exhibiting parallel temporal allele frequency change across replicates (‘adaptive sites’) and candidate loci where we had *a priori* predictions for a contribution to rapid adaptation. These candidate loci were derived from: 1) lists of SNPs and genes previously identified as involved in seasonal adaptation or adaptation over monthly intervals in similar temporal evolution experiments (57, 72) (details in Table S2), 2) loci from genotype–phenotype mapping of fecundity and starvation tolerance (73, 116), and 3) an *a priori* list of candidate genes based on knowledge of the molecular pathways involved in ion regulation in insects (Table S4). To test for the predictability of genomic architecture of adaptation the bedtools-intersect tool was used to examine the degree of overlap between candidate gene intervals ±500bp and all adaptive sites (GCF_000001215.4) (117). The lists of candidate SNPs and those within candidate gene regions were tested for significant hypergeometric overlap with adaptive sites using the *phyper* tool (stats *R*-package). In addition, to test whether entire sets of predictors enabled prediction we conducted one-way non-parametric Wilcoxon rank sum tests using the *wilcox.test* function (stats *R*-package) to compare the overlap of candidates and matched sets of loci to adaptive sites. These tests were run using unadjusted GLM p-values at SNP- and gene-level resolutions. Matched genes were selected randomly from those that have a similar base pair length as candidates (±25%) and located on the same chromosome arm. Rank sum tests at the gene-level were conducted on lists of median p-values summarized per gene. At the SNP-level, sites in each gene pair were sampled to the minimum number of loci between the candidate or matched set gene prior to running rank sum tests. Each of these variations of rank sum tests were run for 1000 iterations of randomized matched set selection. We report the percentage of iterations producing *p*-values < 0.05 and the median *p*-value across all iterations.

## Acknowledgements

We thank August Goldfischer, Ozan Kiratli, and Subhash Rajpurohit for help with experiment procedures. Thank you to members of the Rudman Lab for helpful comments and to Timur Gabidulin for an illustration used in Figure 1.

## Author Contributions

SMR and PSS designed the research in consultation with HAM and PDJ; SMR performed the field experiment; RES and PDJ generated elemental data; SMR, JFH, and SIG analyzed genomic data; SMR and RES analyzed phenotypic data and prepared figures; SMR wrote the paper with input from all authors.

