## Supplemental figures and tables for "Ecological stoichiometry and life history theory, not the identity of genomic variants, predict rapid adaptation"

**This file includes:** Figures S1-S3, Tables S1-S6

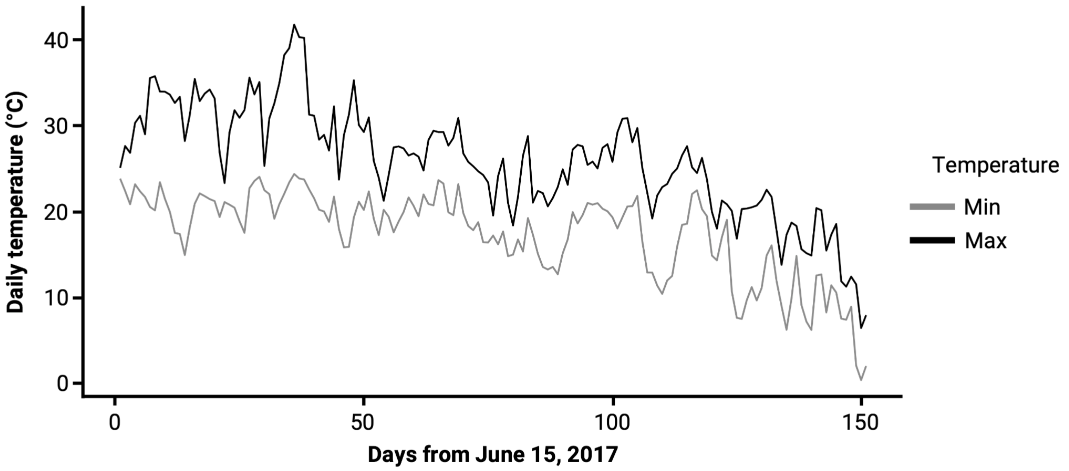


**Figure S1.** The mean daily minimum and maximum temperature (°C) across the eight replicate outdoor cages during the experimental period (June 15 to November 12, 2017).

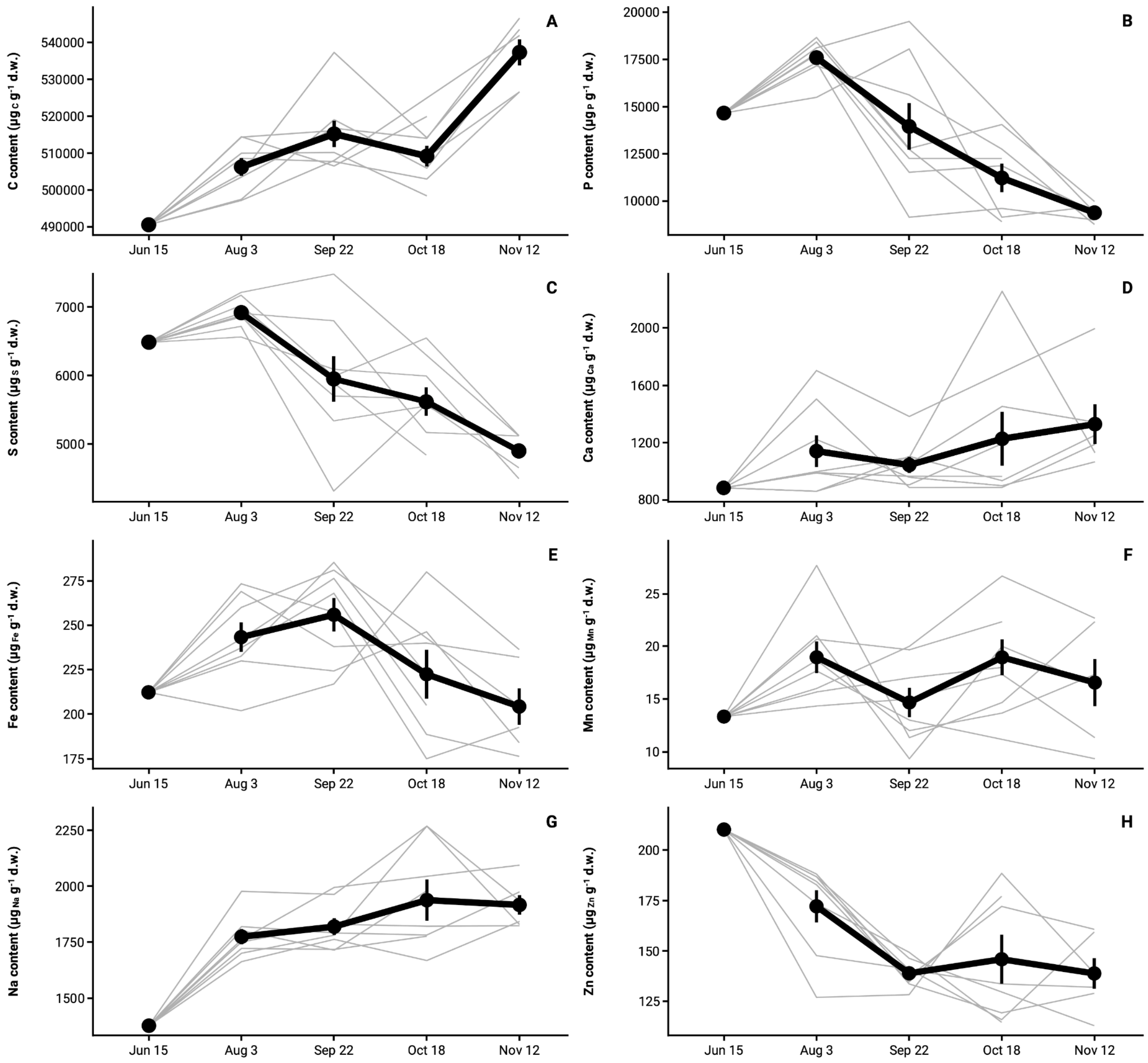


**Figure S2.** Panels A-H show evolution of elemental composition for C, P, S, Ca, Fe, Mn, Na, and Zn, respectively, measured as $\mu g_{element}$ $g^{-1}$ d.w. over the seasonal period. Elemental compositions were measured for pools of individuals from each population following 2 generations of common garden rearing, and hence show genetic change over time in composition. Gray lines represent individual replicate populations and black points (±SE) and lines represent mean values.

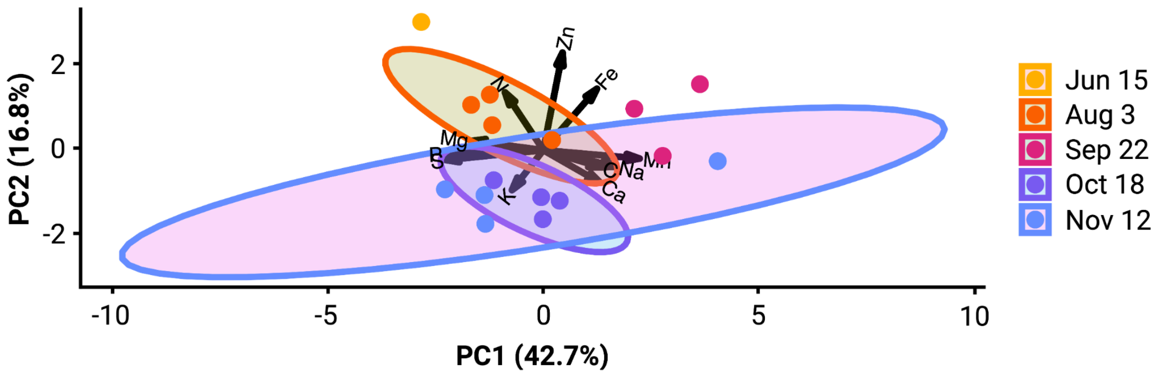


**Figure S3.** The ionomes of *Drosophila melanogaster* founder populations that were held indoors at large population sizes and at constant conditions in the lab. In contrast to field populations, adaptive tracking was not a primary driver of stoichiometric variation (Figure 3E, PC1).

**Table S1 .** Gene Ontology Enrichment table for genes significant in pairwise GLMs at an unadjusted $p < 0.05$. Only the top 20 GO categories are displayed for each time-point contrast, most of which overlap across contrasts. Empty cells may still be enriched within a contrast beyond the top 20 terms.

**Table S2.** Hypergeometric overlap p-values for comparisons between sets of SNPs significant in pairwise time-point contrasts (GLM FDR < 0.05) and candidate SNPs for four phenotypes.

| | | | *SNPs within candidate genes* | | |
|---|---|---|---|---|---|
| *Contrast* | *Ion-transport (N=28,896)* | *Rudman Seasonal (N=32,110)* | *Bitter Seasonal (N=1,271,822)* | *Fecundity (N=7,092)* | *Starvation (N=9,881)* |
| Sep 22 *vs.* Oct 18 (N = 5) | 1 (n=0) | 1 (n=0) | 0.96 (n=2) | 1 (n=0) | 1 (n=0) |
| Oct 18 *vs.* Nov 12 (N=7) | 0.1 (n=1) | 1 (n=0) | 0.59 (n=5) | 1 (n=0) | 1 (n=0) |
| Sep 22 *vs.* Nov 12 (N=13) | 1 (n=0) | 1 (n=0) | 0.97 (n=6) | 1 (n=0) | 1 (n=0) |
| | | | *Candidate SNPs* | | |
| *Contrast* | *Ion-transport* | *Rudman Seasonal (N=47)* | *Bitter Seasonal (N=232,696)* | *Fecundity (N=11)* | *Starvation (N=16)* |
| Sep 22 *vs.* Oct 18 (N = 5) | N/A | 1 (n=0) | 0.48 (n=1) | 1 (n=0) | 1 (n=0) |
| Oct 18 *vs.* Nov 12 (N=7) | N/A | 1 (n=0) | 1 (n=0) | 1 (n=0) | 1 (n=0) |
| Sep 22 *vs.* Nov 12 (N=13) | N/A | 1 (n=0) | 0.82 (n=1) | 1 (n=0) | 1 (n=0) |

**Table S3.** Hypergeometric overlap p-values for comparisons between sets of SNPs significant in pairwise time-point contrasts (GLM unadjusted-$p < 0.01$) and candidate SNPs for four phenotypes.

*SNPs within candidate genes*

| *Contrast* | *Ion-transport (N=28,896)* | *Rudman Seasonal (N=32,110)* | *Bitter Seasonal (N=1,271,822)* | *Fecundity (N=7,092)* | *Starvation (N=9,881)* |
|---|---|---|---|---|---|
| Sep 22 *vs.* Oct 18 (N = 4,319) | 0.93 (n=55) | 0.72 (n=69) | 0.79 (n=2,890) | 0.46 (n=17) | 0.5 (n=23) |
| Oct 18 *vs.* Nov 12 (N=3,979) | 0.47 (n=62) | 0.9 (n=58) | 0.98 (n=2,637) | 0.43 (n=16) | 0.76 (n=18) |
| Sep 22 *vs.* Nov 12 (N=6,855) | 0.56 (n=104) | 0.58 (n=115) | 0.69 (n=4,653) | 0.67 (n=24) | 0.58 (n=35) |

*Candidate SNPs*

| *Contrast* | *Ion-transport* | *Rudman Seasonal (N=47)* | *Bitter Seasonal (N=232,696)* | *Fecundity (N=11)* | *Starvation (N=16)* |
|---|---|---|---|---|---|
| Sep 22 *vs.* Oct 18 (N = 4,319) | N/A | 1 (n=0) | 0.08 (n=602) | 1 (n=0) | 1 (n=0) |
| Oct 18 *vs.* Nov 12 (N=3,979) | N/A | 1 (n=0) | 0.12 (n=526) | 1 (n=0) | 1 (n=0) |
| Sep 22 *vs.* Nov 12 (N=6,855) | N/A | 1 (n=0) | **0.03** (n=954) | 1 (n=0) | 1 (n=0) |

**Table S4.** List of candidate genes known to be involved in ion-transport. List was established as an *a prior* prediction based on expert knowledge of insect ion transport physiology (*H. MacMillan*, ) prior to conducting genomic sequencing and analyses.

| *Gene* | *Name* | *Reason* |
|---|---|---|
| alpha-Cat | alpha catenin | septate junction component |
| aPKC | DaPKC | septate junction component |
| atpalpha | Na pump alpha subunit | alpha subunit of sodium pump |
| baz | Bazooka | septate junction component |
| bou | Boudin | septate junction component |
| Ca-Ma2d | Ca2+ channel Muscle-specific alpha2/delta subunit | calcium channel |
| Ca-alpha1D | Ca2+-channel protein α1 subunit D | calcium |
| Ca-alpha1T | Ca2+-channel protein α1 subunit T | calcium |
| Ca-beta | Ca2+-channel-protein-β-subunit | calcium |
| cac | cacophony | calcium channel |
| capa | Capability | neuropeptide involved in ion balance |
| capaR | Capa receptor | receptor for neuropeptide involved in ion balance |
| pck | pickel | septate junction component |
| dlg1 | Discs large | septate junction component |
| cold | coiled | septate junction component |
| mesh | mesh | septate junction component |
| arm | Armadillo | septate junction component |
| bark | Bark beetle | septate |
| cno | Canoe | septate junction component |
| cora | coracle | septate junction component |
| crb | Crumbs | septate junction component |
| crim | Crimpled | septate junction component |
| crok | Crooked | septate junction component |
| Dh31 | Diuretic hormone 31 | neuropeptide involved in ion balance |

| | | |
|---|---|---|
| Dh31-R | DH31 receptor 1 | receptor |
| Dh44 | Diuretic hormone 44 | neuropeptide involved in ion balance |
| Dh44-R1 | Dh44 receptor 1 | receptor for neuropeptide involved in ion balance |
| Dh44-R2 | DH 44 receptor 2 | receptor for neuropeptide involved in ion balance |
| Drip | Drip | putative aquaporin (water balance is tied to ion balance) |
| eag | ether a go-go | Potassium channel |
| Eglp1 | Entomoglyceroporin 1 | putative aquaporin (water balance is tied to ion balance) |
| Eglp2 | Entomoglyceroporin 2 | putative aquaporin (water balance is tied to ion balance) |
| Eglp3 | Entomoglyceroporin 3 | putative aquaporin (water balance is tied to ion balance) |
| Eglp4 | Entomoglyceroporin 4 | putative aquaporin (water balance is tied to ion balance) |
| Elk | Eag-like K+ channel | potassium channel |
| Fas3 | Fasciclin III | septate junction component |
| Gli | Gliotactin | septate junction component |
| Hk | Hyperkinetic | potassium channel |
| Irk1 | Inwardly rectifying potassium channel 1 | Potassium channel |
| Irk2 | Inwardly rectifying potassium channel 2 | Potassium channel |
| Irk3 | Inwardly rectifying potassium channel 3 | potassium channel |
| ITP | Ion transport peptide | neuropeptide involved in ion balance |
| KCNQ | KCNQ potassium channel | potassium channel |
| kune | kune-kune | septate junction component |
| l(2)gl | Lgl | septate junction component |
| Lac | Lachesin | septate junction component |
| Lk | Leucokinin | neuropeptide involved in ion balance |
| Lkr | Leucokinin receptor | receptor for neuropeptide involved in ion balance |

| | | |
|---|---|---|
| Nckx30C | Nckx30C | calcium-potassium:sodium exchanger |
| Nha2 | Na+/H+ antiporter 2 | Sodium-hydrogen exchanger |
| Nhe1 | Na+/H+ exchanger 1 | Sodium-hydrogen exchanger |
| NHE3 | Na+/H+ exchanger 3 | Sodium-hydrogen exchanger |
| NKAIN | Na,K-ATPase Interacting | regulator of sodium pump |
| NKCC | sodium potassium chloride cotransporter | exchanges ions across membranes |
| Nrg | Neuroglian | septate junction component |
| nrv1 | nervana 1 | Beta subunit of sodium pump |
| nrv2 | Nrv2 | septate junction component |
| nrv2 | nervana 2 | beta subunit of sodium pump |
| nrv3 | nervana 3 | Beta subunit of sodium pump |
| Nrx-IV | Neurexin IV | septate junction component |
| Ork1 | Open rectifier K+ channel 1 | Potassium channel |
| par-6 | DmPar6 | septate junction component |
| para | paralytic | sodium channel alpha subunit |
| PMCA | plasma membrane calcium ATPase | calcium pump |
| Prip | Prip | putative aquaporin (water balance is tied to ion balance) |
| scrib | Scribble | septate junction component |
| sdt | Stardust | septate junction component |
| sei | Seizure | potassium channel |
| SERCA | Sarco/endoplasmic reticulum Ca(2+)-ATPase | Calcium pump |
| Sh | Shaker | potassium channel |
| Shab | Shaker conjugate b | Potassium channel |
| Shal | Shaker conjugate 1 | Potassium channel |
| Shaw | Shaker conjugate w | Potassium channel |
| shg | Shotgun | septate junction component |
| sinu | sinuous | septate junction component |
| Slo | Slowpoke | Potassium channel |

| | | |
|---|---|---|
| Ssk | Snakeskin | septate junction component |
| Teh1 | tipE homolog 1 | sodium channel beta subunit |
| Teh2 | tipE homolog 2 (a.k.a. Vacuolar H+ ATPase M9.7 subunit c) | sodium channel beta subunit |
| Teh3 | tipE homolog 3 | sodium channel beta subunit |
| Teh4 | tipE homolog 4 | sodium channel beta subunit |
| tipE | temperature-induced paralytic E | sodium channel beta subunit |
| Tsf2 | Melanotransferin/Transferin 2 | septate junction component |
| vari | Varicose | septate junction component |
| Vha100-1 | Vacuolar H+ ATPase 100kD subunit 1 | component of V-ATPase (creates H+ gradients used to drive other ions across membranes) |
| Vha100-2 | Vacuolar H+ ATPase 100kD subunit 2 | component of V-ATPase (creates H+ gradients used to drive other ions across membranes) |
| Vha100-3 | Vacuolar H+ ATPase 100kD subunit 3 | component of V-ATPase (creates H+ gradients used to drive other ions across membranes) |
| Vha100-4 | Vacuolar H+ ATPase 100kD subunit 4 | component of V-ATPase (creates H+ gradients used to drive other ions across membranes) |
| Vha100-5 | Vacuolar H+ ATPase 100kD subunit 5 | component of V-ATPase (creates H+ gradients used to drive other ions across membranes) |
| Vha13 | Vacuolar H+ ATPase 13kD subunit | component of V-ATPase (creates H+ gradients used to drive other ions across membranes) |
| Vha14-1 | Vacuolar H+ ATPase 14kD subunit 1 | component of V-ATPase (creates H+ gradients used to drive other ions across membranes) |
| Vha14-2 | Vacuolar H+ ATPase 14kD subunit 2 | component of V-ATPase (creates H+ gradients used to drive other ions across membranes) |

| | | |
|---|---|---|
| Vha16-1 | Vacuolar H+ ATPase 16kD subunit 1 | component of V-ATPase (creates H+ gradients used to drive other ions across membranes) |
| Vha16-2 | Vacuolar H+ ATPase 16kD subunit 2 | component of V-ATPase (creates H+ gradients used to drive other ions across membranes) |
| Vha16-3 | Vacuolar H+ ATPase 16kD subunit 3 | component of V-ATPase (creates H+ gradients used to drive other ions across membranes) |
| Vha16-4 | Vacuolar H+ ATPase 16kD subunit 4 | component of V-ATPase (creates H+ gradients used to drive other ions across membranes) |
| Vha16-5 | Vacuolar H+ ATPase 16kD subunit 5 | component of V-ATPase (creates H+ gradients used to drive other ions across membranes) |
| Vha26 | Vacuolar H+ ATPase 26kD subunit | component of V-ATPase (creates H+ gradients used to drive other ions across membranes) |
| Vha36-1 | Vacuolar H+ ATPase 36kD subunit 1 | component of V-ATPase (creates H+ gradients used to drive other ions across membranes) |
| Vha36-2 | Vacuolar H+ ATPase 36kD subunit 2 | component of V-ATPase (creates H+ gradients used to drive other ions across membranes) |
| Vha36-3 | Vacuolar H+ ATPase 36kD subunit 3 | component of V-ATPase (creates H+ gradients used to drive other ions across membranes) |
| Vha44 | Vacuolar H+ ATPase 44kD subunit | component of V-ATPase (creates H+ gradients used to drive other ions across membranes) |
| Vha55 | Vacuolar H+-ATPase 55kD subunit | component of V-ATPase (creates H+ gradients used to drive other ions across membranes) |
| Vha68-1 | Vacuolar H+ ATPase 68kD subunit 1 | component of V-ATPase (creates H+ gradients used to drive other ions across membranes) |

| | | |
|---|---|---|
| Vha68-2 | Vacuolar H+ ATPase 68 kDa subunit 2 | component of V-ATPase (creates H+ gradients used to drive other ions across membranes) |
| Vha68-3 | Vacuolar H+ ATPase 68kD subunit 3 | component of V-ATPase (creates H+ gradients used to drive other ions across membranes) |
| VhaAC39-1 | Vacuolar H+ ATPase AC39 subunit 1 | component of V-ATPase (creates H+ gradients used to drive other ions across membranes) |
| VhaAC39-2 | Vacuolar H+ ATPase AC39 subunit 2 | component of V-ATPase (creates H+ gradients used to drive other ions across membranes) |
| VhaM9.7-a | Vacuolar H+ ATPase M9.7 subunit a | component of V-ATPase (creates H+ gradients used to drive other ions across membranes) |
| VhaM9.7-b | Vacuolar H+ ATPase M9.7 subunit b | component of V-ATPase (creates H+ gradients used to drive other ions across membranes) |
| VhaM9.7-d | Vacuolar H+ ATPase M9.7 subunit d | component of V-ATPase (creates H+ gradients used to drive other ions across membranes) |
| VhaPPA1-1 | Vacuolar H+ ATPase PPA1 subunit 1 | component of V-ATPase (creates H+ gradients used to drive other ions across membranes) |
| VhaPPA1-2 | Vacuolar H+ ATPase PPA1 subunit 2 | component of V-ATPase (creates H+ gradients used to drive other ions across membranes) |
| VhaSFD | Vacuolar H+-ATPase SFD subunit | component of V-ATPase (creates H+ gradients used to drive other ions across membranes) |
| zyd | Zydeco | sodium calcium exchanger |

**Table S5.** Median *p*-values for 1000 iterations of WilcoxonRank Sum tests of candidate SNPs *vs.* matched SNP sets.

*SNPs within candidate genes*

| *Contrast* | *Ion-transport (N=28,896)* | *Rudman Seasonal (N=32,110)* | *Fecundity (N=7,092)* | *Starvation (N=9,881)* |
|---|---|---|---|---|
| Sep 22 *vs.* Oct 18 | 0.74 | **7.1E-04** | 0.80 | 0.09 |
| Oct 18 *vs.* Nov 12 | **0.02** | 0.36 | 0.79 | 0.42 |
| Sep 22 *vs.* Nov 12 | 0.64 | 0.88 | 0.28 | **0.01** |

*Candidate SNPs*

| *Contrast* | *Ion-transport* | *Rudman Seasonal (N=47)* | *Fecundity (N=11)* | *Starvation (N=16)* |
|---|---|---|---|---|
| Sep 22 *vs.* Oct 18 | N/A | 0.39 | 0.94 | 0.61 |
| Oct 18 *vs.* Nov 12 | N/A | 0.12 | 0.43 | 0.80 |
| Sep 22 *vs.* Nov 12 | N/A | 0.67 | 0.57 | 0.48 |

**Table S6.** Percent of *p*-values < 0.05 for 1000 iterations of WilcoxonRank Sum tests for candidate SNPs *vs.* matched SNP sets.

| | | *SNPs within candidate genes* | | |
|---|---|---|---|---|
| *Contrast* | *Ion-transport (N=28,896)* | *Rudman Seasonal (N=32,110)* | *Fecundity (N=7,092)* | *Starvation (N=9,881)* |
| Sep 22 *vs.* Oct 18 | 0.9 | **88.3** | 0.8 | 39.6 |
| Oct 18 *vs.* Nov 12 | **64.2** | 11 | 1.9 | 8.9 |
| Sep 22 *vs.* Nov 12 | 4.6 | 1.8 | 20.6 | **67.6** |
| | | *Candidate SNPs* | | |
| *Contrast* | *Ion-transport* | *Rudman Seasonal (N=47)* | *Fecundity (N=11)* | *Starvation (N=16)* |
| Sep 22 *vs.* Oct 18 | NA | 1.6 | 0 | 0.2 |
| Oct 18 *vs.* Nov 12 | NA | 25.2 | 1.7 | 0 |
| Sep 22 *vs.* Nov 12 | NA | 0.2 | 0.8 | 2.9 |
